# Nailpolish: Reference-free UMI deduplication and consensus read generation

**DOI:** 10.64898/2026.09.25.754331

**Authors:** Oliver Y. Cheng, Jia Wei Tan, Nadia M. Davidson

## Abstract

Long-read sequencing captures full-length transcripts but suffers elevated error rates, and PCR duplicates can distort quantification unless accounted for with UMIs. We present Nailpolish, a fast reference-free tool that error-corrects and deduplicates UMI-tagged long reads by generating consensus sequences via partial order alignment, while separating false duplicates arising from UMI collisions. Across eleven Oxford Nanopore Technologies and Pacific Biosciences datasets spanning bulk, single-cell, spatial, and targeted protocols, Nailpolish reduced per-read error rates and outperformed existing deduplication tools.

## Background

Long-read sequencing is transforming transcriptomics by capturing full-length isoforms, alternative splicing and fusion transcripts that short reads can only infer. Paired with single-cell and spatial technologies, it reveals the complexity of the transcriptome at a new resolution (1). One issue that remains underappreciated in long-read analysis, however, is the presence of PCR duplicates (**Figure 1A**), which can distort quantification if not accounted for. This is well recognised in single-cell transcriptomics, where it is addressed with unique molecular identifiers (UMIs): short sequences incorporated into the cDNA during library preparation, prior to PCR amplification (2). As UMIs tag each original molecule, they enable the computational deduplication of PCR duplicates. In short-read bulk sequencing, duplicates can instead be collapsed by coordinate, since they share start and end positions after genomic alignment (3). However, this strategy is unreliable for long-read data. Therefore, UMIs are increasingly being used for bulk sample deduplication.

**Figure 1:**
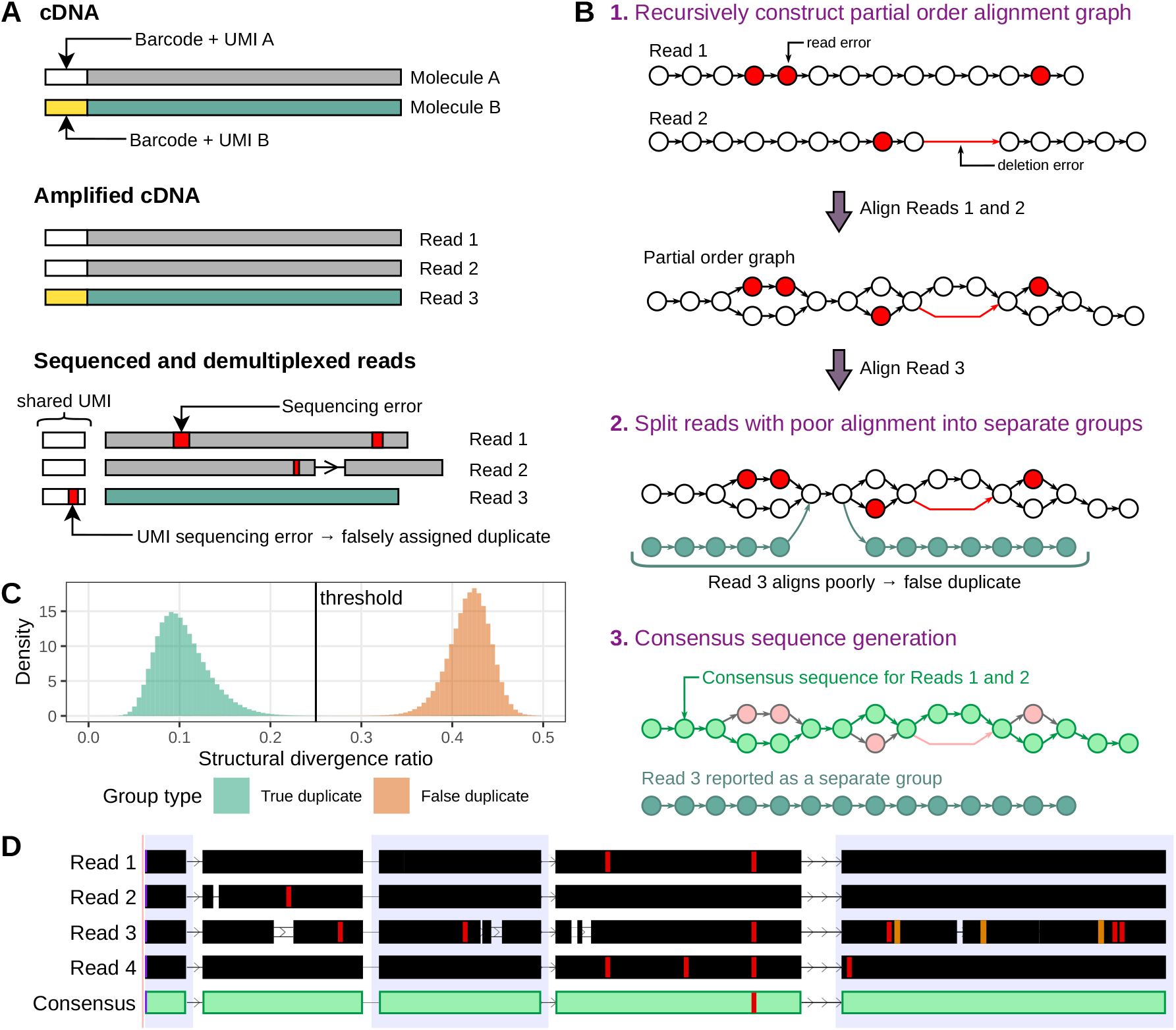
The Nailpolish consensus calling and deduplication approach. **A)** Each cDNA molecule is tagged with an identifier, such as a cell barcode and UMI, prior to amplification, so that reads with the same identifier can be deduplicated. Sequencing errors occur across the reads, including in the UMI, which may cause reads to falsely be classified as duplicates. **B)** Nailpolish collects all reads which share the same identifier, then aligns each read into a partial order alignment graph. Read alignments with a ‘structural divergence ratio’ beyond a set threshold are deemed ‘false duplicates,’ reverted, and assigned to a new group. A consensus sequence is then derived from each subgroup. **C)** Distribution of the structural divergence ratio for true and false duplicates derived from real data, with Nailpolish’s chosen threshold value. **D)** Real example of a duplicate group (black tracks) and its Nailpolish consensus sequence (green track). Single base mismatches (red), insertions (orange) and deletions (gaps) with respect to the reference genome are shown. Data were taken from ONT RNA-Seq from You et al. (17) and read alignment visualisation adapted from the UCSC Genome Browser (18).

A range of computational tools exist for UMI-based deduplication, but they vary in the sequencing protocols they support, whether they require prior alignment to a reference, and importantly, how they select a representative read per duplicate group (**Supplementary Table 1**). One of the most widely used tools is UMI-tools (4), which was originally designed for short-read data. It accepts an aligned BAM file and generates a new BAM retaining a representative read per duplicate group. While efficient, this approach does not exploit the information available to construct a consensus sequence in which errors are resolved. This is a notable limitation for platforms such as those from Oxford Nanopore Technologies (ONT) (5), which are subject to elevated error rates. There are alternative tools which generate consensus sequences (6–9); however, the majority of these have also been developed for short-read data and rely on per-base voting, making them poorly suited to correct the insertion- and deletion-heavy errors found in long-read data. Long-read specific methods have been developed, with a prime example being ONT’s duplex basecalling via Dorado (10), but this works on a single molecule and not PCR duplicates. Other tools require specialised library preparation such as the addition of bespoke adapters and UMIs (11,12). Some tools do exist which could be used on standard long-read sequencing data, such as SiCeLoRe (13,14). However, these tools require preprocessing from upstream steps in their pipelines, and reads must first be aligned to a reference genome or transcriptome, limiting their utility in non-model species or fusion gene detection in cancer. Ultimately, existing methods either work on limited library types, are not appropriate for the error profile of long-read data, or cannot be applied prior to reference alignment.

## Results and discussion

To address the limitations of existing methods, we developed Nailpolish, a reference-free tool for error correction of PCR duplicates in UMI-tagged long-read sequencing data. When given UMI-demultiplexed reads in FASTQ format, Nailpolish produces duplicate statistics for quality assessment and generates a consensus FASTQ from duplicate groups (**Figure 1B-D**). Nailpolish is distributed as a single executable binary written in Rust and is available at https://github.com/davidsongroup/nailpolish.

Nailpolish only requires post-demultiplexed read data, making it compatible with any type of data where UMIs can be extracted. By default, Nailpolish assumes the barcode and UMI encoding produced by Flexiplex (15) and BLAZE (16), but this is modifiable through a user-provided regular expression or from a set of presets. Identifiers may take a variety of forms: a standalone UMI, as in bulk protocols; a barcode paired with a UMI, as in droplet-based single-cell sequencing protocols; or an arbitrary number of UMIs, as seen in combinatorial barcoding strategies such as split-seq and 10x Visium HD. For brevity, we refer to all these composite identifiers as UMIs.

To improve computational efficiency, Nailpolish first indexes input files to construct a lookup table serialised on disk, allowing random-access read retrieval. Both uncompressed and gzip-compressed input are supported. Indexation is fast and memory efficient, completing in under 10 minutes with one thread on all datasets tested. An optional HTML summary page can be produced after indexation with estimated duplicate distributions and sequencing saturation (**Supplementary Figure 1**).

For consensus calling, reads sharing identical UMIs are collected into duplicate groups. Each group is further partitioned into *subgroups*, each holding a partial order alignment graph. Every group initially contains a single subgroup. Candidate reads are added to the subgroup in descending order of their mean Phred quality score, with sequence alignment performed using the partial order alignment algorithm (19) via native bindings to the SPOA library (20) (**Figure 1B**). To detect false duplicates arising from UMI collisions, every candidate alignment is evaluated before being inserted into the graph. False duplicates introduce many new nodes along their length. We quantify this using a structural divergence ratio, defined as the ratio of newly added nodes to the aligned length of the candidate read within the graph. If the ratio exceeds a threshold (default 0.25), the alignment is discarded and the read tested against the next subgroup. A read that does not align with any subgroup is placed into a new subgroup. The structural divergence ratio threshold was determined on a labelled set of true and false duplicates derived from real data (**Figure 1C, Methods**).

By default, reads exceeding 15,000bp, or belonging to duplicate groups with more than 250 reads, are output unchanged. These thresholds are user-configurable and prevent erroneous reads from having an outsized effect on runtime. Alternatively, large duplicate groups can be subsampled to the maximum group size prior to consensus calling, either pseudorandomly or by selecting the N-longest reads.

After all reads in a group have been assigned to a subgroup, a consensus sequence is derived from each using the heaviest bundle algorithm defined by Lee (21) and implemented in SPOA. Identifying metadata such as the group size, barcode, and UMI are appended to the output read header as FASTQ comments (allowing the identifiers to be incorporated as SAM tags after alignment using the -y parameter of minimap2 (22)). As SPOA produces a consensus sequence without base qualities, the quality score for each base is set to the highest Phred score across all reads which support the consensus base at that position.

Nailpolish’s performance was evaluated on diverse long-read datasets spanning ONT and PacBio platforms across multiple protocols including bulk, single-cell, spatial, targeted, and untargeted sequencing (**Figure 2A, Supplementary Table 2**). Across these datasets we found duplication rates were lowest for long-read single-cell protocols but significant in bulk, spatial and targeted protocols, motivating the need for a robust and flexible deduplication tool (**Figure 2B**).

**Figure 2:**
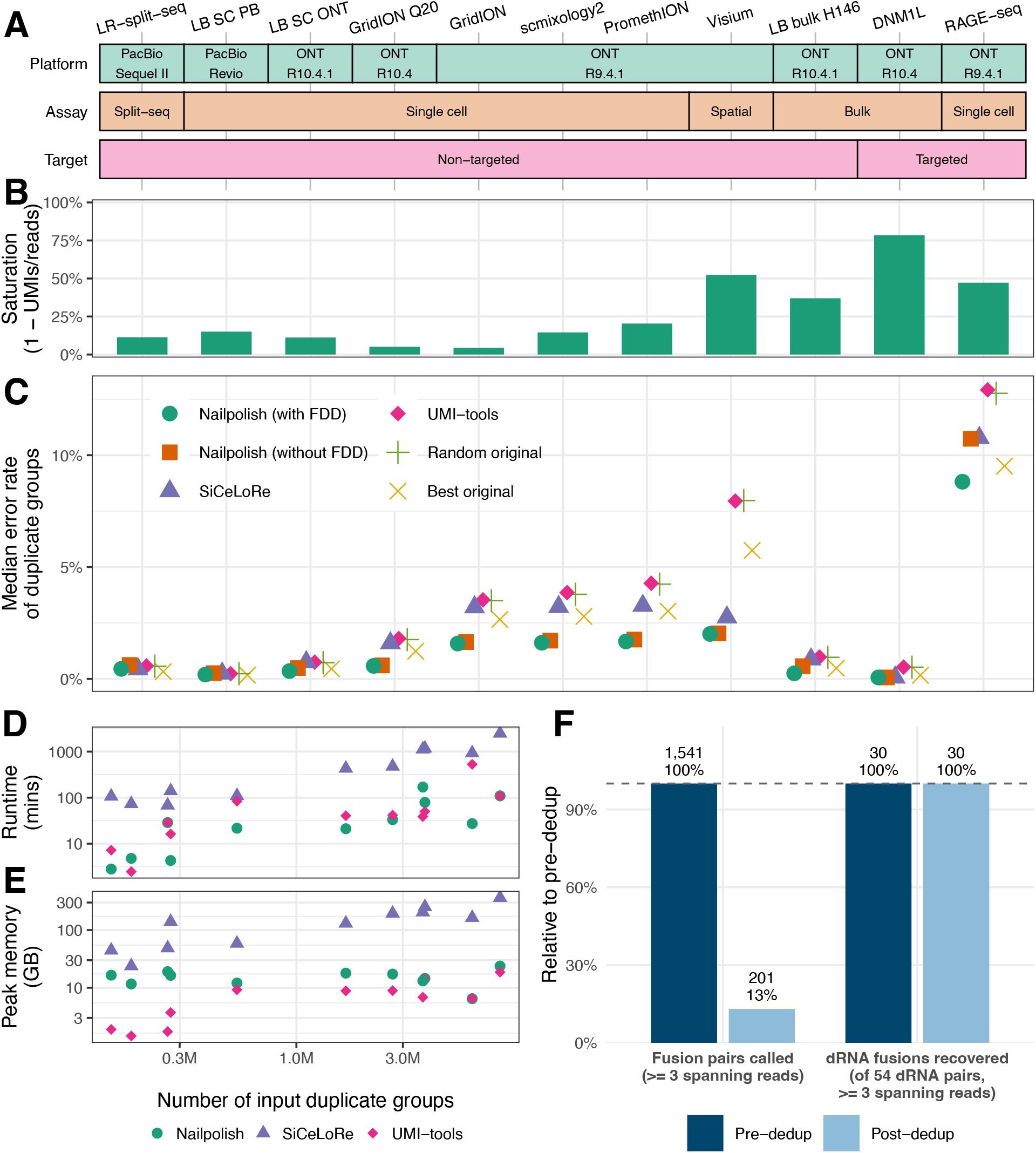
Performance of Nailpolish. **A)** The platform, assay, and target enrichment for each dataset tested. **(B)**Library saturation, computed as 1 - (number of unique UMIs/ total reads). **C)** Median error rate of representative reads across duplicate groups. The representative read for each group was selected using: Nailpolish consensus with and without false duplicate detection (FDD); SiCeLoRe consensus; UMI-tools representative read selection; random selection from the group; and the read with the lowest ground-truth error rate. **D)** Wall-clock runtime (log scale) of deduplication tools Nailpolish, SiCeLoRe, and UMI-tools plotted against the number of input duplicate groups. Nailpolish was run using 16 threads; UMI-tools with its maximum of 1 thread and SiCeLoRe with 16 threads. **E)** The peak memory (log scale) of deduplication tools Nailpolish, SiCeLoRe, and UMI-tools plotted against the number of input duplicate groups. **F)** The number of high-confidence gene pairs called by JAFFAL at ≥3 spanning reads for the H146 bulk ONT cDNA sample before and after Nailpolish deduplication (left) and how many of the 54 ONT direct-RNA truth pairs those calls recover (right).

To assess error correction performance, reads were aligned to their respective reference genomes using minimap2. Per-read error rate was defined as the ratio of alignment edit distance (as reported by minimap2) to alignment length (**Methods**). For each duplicated group, we compared three approaches to producing a representative read: random selection, the read with the lowest error rate (as defined above) and consensus calling with Nailpolish. Nailpolish consensus reads had the lowest median error rates in 9 of the 11 datasets (**Figure 2C**). In the remaining 2 datasets (both PacBio), the error rate was already low (0.3% and 0.2%), and Nailpolish’s error rate only marginally higher than selecting the lowest error read. A limitation of Nailpolish is that consensus calling assumes errors are independent across duplicate reads. Systematic homopolymer errors are common in data from ONT R9.4.1 or earlier flow cells (23) and may explain why error rates remain >1% in these datasets even after error correction.

To assess the impact of Nailpolish’s false duplicate detection (FDD), we compared it against Nailpolish with it disabled. FDD reduced the estimated sequencing saturation in all datasets, with reductions ranging from 0.20% to 12% (**Supplementary Figure 2**), and provided a modest drop in median error rate (**Figure 2C**). This demonstrates the importance of accounting for UMI collisions. Nailpolish effectively resolves these without relying on a reference alignment, which other tools use to distinguish falsely grouped reads by their gene of origin.

Next, we evaluated Nailpolish against alternative deduplication tools: UMI-tools’ dedup command, which selects a representative read per duplicate group, and SiCeLoRe’s ComputeConsensus pipeline, which produces a consensus read per group. All tools were provided with barcodes and UMIs demultiplexed by Flexiplex. UMI-tools and SiCeLoRe were provided with mapped reads, whereas Nailpolish operated directly on unmapped reads. UMI-tools produced reads with an error rate similar to selecting a random read per duplicate group, and as a consequence had a higher error rate than Nailpolish across all datasets tested. Compared to SiCeLoRe, Nailpolish produced lower median error rates across 9 out of 11 datasets.

Computational time and memory were evaluated for each tool on a Linux high-performance computer. Both Nailpolish and UMI-tools were fast and efficient, completing in a comparable amount of time that never exceeded 8 hours or 24 GB on all datasets (**Figure 2D-E**). Nailpolish’s similar run-time was a consequence of its multi-threading (16 threads) compared to UMI-tools, which could only be run using 1 thread. SiCeLoRe was found to have considerably higher resource requirements, taking up to 41 hours with 16 threads and needing up to 360 GB of RAM.

Finally, we examined the downstream effects of deduplication, taking an example use case of single nucleotide variant (SNV) and fusion calling, on the bulk ONT cDNA data from the H146 cancer cell line. The effect on fusion calling was large: the number of distinct fusions called by JAFFAL (24) and supported by at least three reads dropped significantly (**Figure 2F**) after deduplication, likely because trans-splicing background events, usually supported by a single mRNA molecule (24), had their supporting counts inflated by PCR duplication. However, the number of fusion calls shared with matched ONT direct RNA data, which is free from PCR amplification, was unchanged after deduplication, suggesting that deduplication removes false fusion calls without sacrificing true positives. Notably, most long-read fusion callers require unmapped reads, demonstrating the necessity of reference-free deduplication methods. For SNVs, we observed a similar but modest trend, where the total number of variants called fell by 13-20% after deduplication, while recovery of variants seen in whole-exome data was similar (**Supplementary Figure 4**). These analyses did not account for missed duplicates due to sequencing errors in UMIs. Therefore, coupling Nailpolish with a reference-free UMI correction tool such as UMI-nea (25) could improve precision in applications such as variant and fusion detection, even further.

## Conclusion

We present Nailpolish, a fast, reference-free tool for error correction and deduplication of UMI-tagged long reads that builds a consensus sequence per duplicate group and separates false duplicates arising from UMI collisions without requiring prior alignment. Across eleven ONT and PacBio datasets spanning bulk, single-cell, spatial, and targeted protocols, Nailpolish reduced per-read error rates relative to UMI-tools and SiCeLoRe while remaining computationally light. Deduplication with Nailpolish was found to enrich true positives in a use case of variant and fusion calling on bulk long-read data, indicating that improvement in read accuracy and quantification translates into more reliable downstream results. Nailpolish requires only demultiplexed FASTQ input, making it straightforward to insert into any UMI-tagged long-read workflow.

## Methods

### Identification and consensus calling of duplicated reads

Nailpolish expects reads in FASTQ format with barcode and UMI information extracted and placed into the header or alternatively a file mapping read IDs to duplicate clusters. By default Nailpolish searches for barcodes and UMIs stored in the FASTQ ‘comment’ format similar to SAM tags, such as CB:Z:<barcode> and UB:Z:<umi>. Other presets are available, such as the format produced by Flexiplex and BLAZE (@<bc>_<umi>). Alternatively, using the Rust crate *regex*, custom header formats are supported.

During the indexing step, Nailpolish scans through the input file and stores the barcode, UMI, and read byte position in an index file. If the input file is compressed, a set of gzip block access points is also created using the *indexed_deflate* crate, enabling random access.

During consensus calling, duplicate groups are read from the source file at locations stored in the index. Non-duplicate groups (with 1 read each) are replicated to the output FASTQ unchanged. For each duplicate group, the reads are first sorted by descending mean Phred quality score. The highest-quality read is used to create the first subgroup and its corresponding partial order alignment (POA) graph. For each following candidate read, alignment first occurs against the first subgroup using the *spoa* library with the default penalty matrix in semi-global alignment mode. If the candidate alignment satisfies the criteria for a true duplicate, the read is incorporated into the subgroup’s alignment graph. Otherwise, this candidate read is considered a false duplicate and tested against each subsequent subgroup. If the candidate read is not considered a true duplicate of any subgroup, it is placed into a new subgroup.

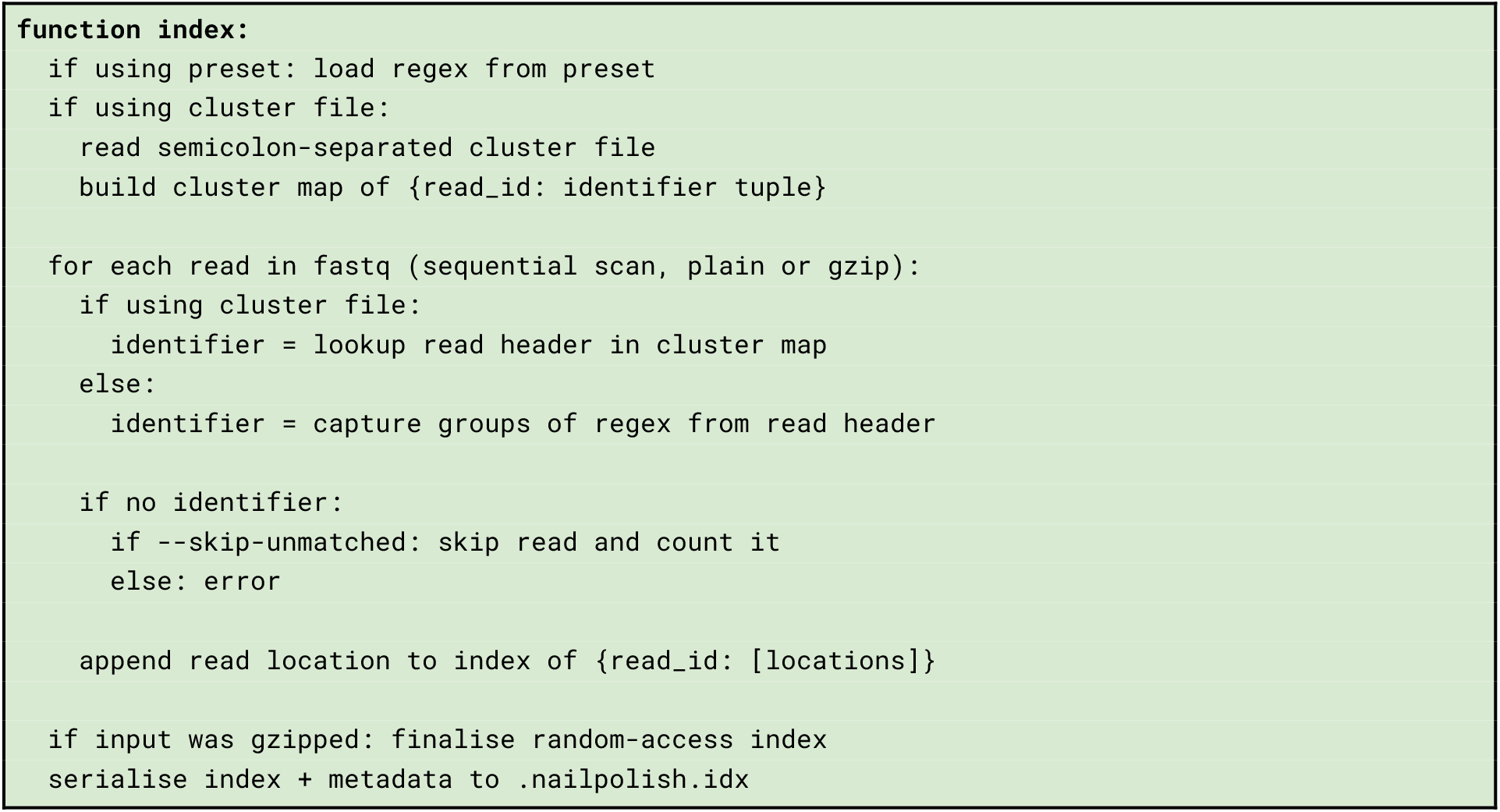

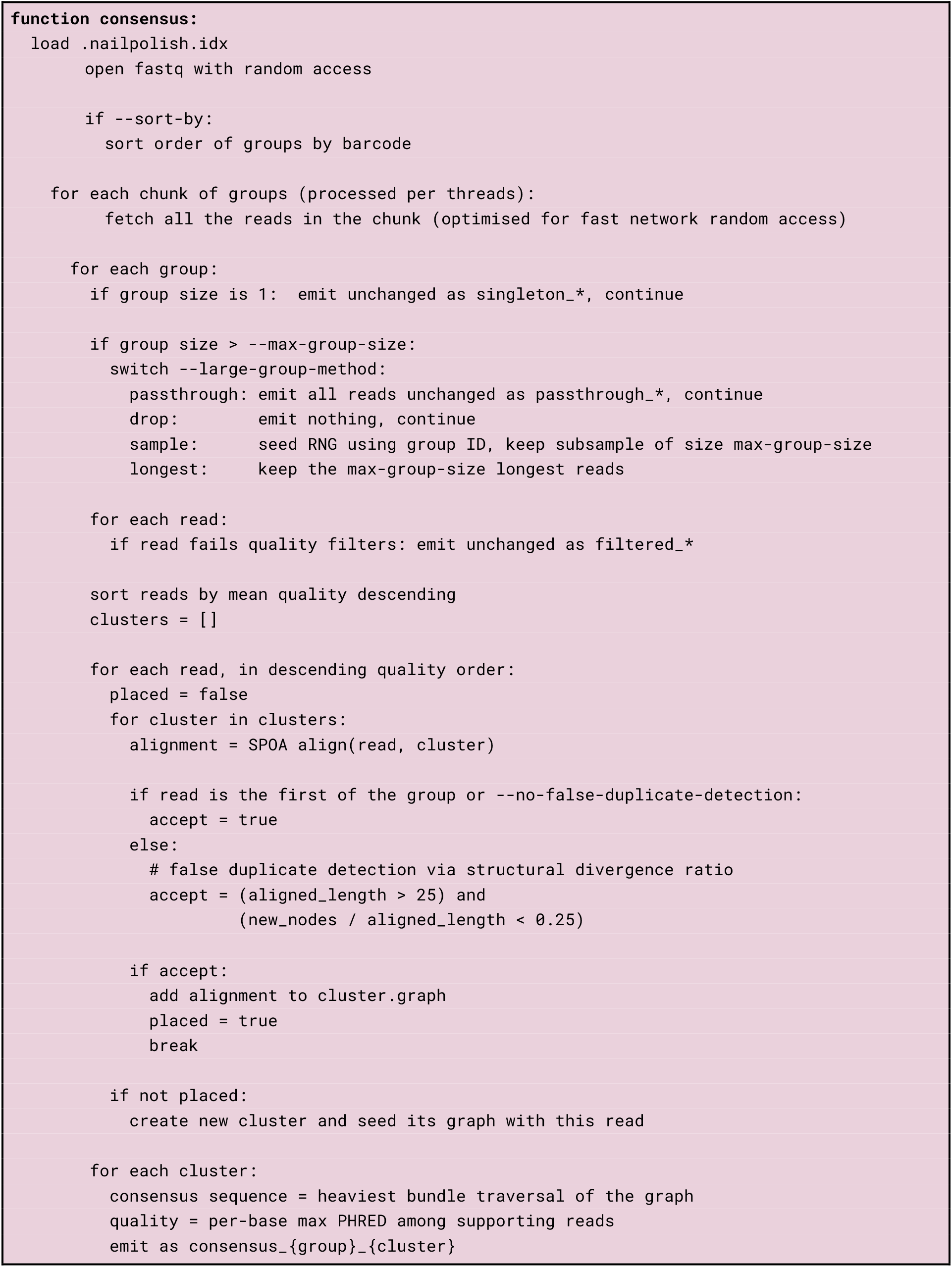

### Clustering of duplicated reads

Alignment of a candidate read against the alignment graph is performed using an embedded build of the spoa library, modified to output the number of new nodes required and existing nodes shared for each alignment. Using this extra data, a structural divergence ratio is computed using the formula

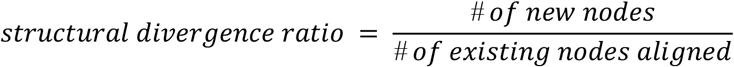

Reads with aligned nodal count under 25 bases, or with a structural divergence ratio below 0.25 (by default) are determined to have failed consensus assembly.

The default structural divergence ratio threshold was determined on a labelled set of true and false duplicates derived from the *Visium* dataset. Reads were aligned to the reference genome with *minimap2*, and read pairs sharing their UMI and aligned chromosome were labelled as ‘true duplicates.’ Synthetic false duplicates were generated from randomly selected read pairs with different UMIs and aligned chromosomes, by assigning both reads an identical artificial UMI. The structural divergence ratio was computed for each pair and an empirical threshold for false duplicates determined to be 0.25 (see Figure 1C). These parameters separated the two distributions with a precision of 0.998, recall of 0.992, and F1 score of 0.995, with misclassifications favouring splitting across subgroups. We subsequently verified our default threshold of 0.25 on all datasets (Supplementary Figure 3).

### Benchmarking

Reads were first demultiplexed using Flexiplex v1.02.5, where the extracted barcodes and UMIs were inserted into the read header as CB:Z:<barcode> UB:Z:<umi>. Flexiplex parameters can be found at https://github.com/davidsongroup/nailpolish_benchmark. For LR-split-seq, the three combinatorial barcodes were concatenated and placed in the CB field. Consensus reads were generated using nailpolish v0.2.2 (commit #6d97184). Indexing was performed with the flags --barcode-regex ‘\tCB:Z:(?<CB>[ATCGNX]+).*\tUB:Z:(?<UB>[ATCGNX]+)’. Consensus generation was performed with the flag -t 16 for performance testing, -t 16 --report-original-reads --extra-stats for comparisons between original and consensus reads, and -t 16 --no-false-duplicate-detection --report-original-reads --extra-stats to assess the performance of Nailpolish without false duplicate detection. Furthermore, for the RAGE-seq dataset, which is targeted and was found to contain very large duplicate groups, the -- large-group-method sample parameter was passed in to all instances of Nailpolish.

To compare against SiCeLoRe and UMI-tools, we ran each tool with the following parameters.

#### SiCeLoRe

Reads were first mapped to the genome reference with minimap2 using -ax splice --secondary=no, and the complete read sequence was set to the CS tag. The original demultiplexed barcode and UMI were retained in the UB and CB tag. Next, spoa v4.1.5 (commit #167bdcc) was downloaded from GitHub and built using GCC v11.5.0. Finally, SiCeLoRe v2.1 (commit #33209a0) was run using OpenJDK v17.0.19:

~~~
java -jar -Xmx400g sicelore-2.1/Jar/Sicelore-2.1.jar ComputeConsensus \
  I={input_bam} O={output_fastq} T=16 \
  CELLTAG=CB UMITAG=UB CDNATAG=CS \
  TMPDIR=“$TMPDIR”
~~~

#### UMI-tools

Reads were first mapped to the genome reference with minimap2 using -ax splice --secondary=no, and the output was coordinate-sorted and indexed using SAMtools. Gene annotations were taken from the GENCODE human (GRCh38, v49) and mouse (GRCm38, M11) comprehensive annotations. Records of type ‘gene’ within chromosomes 1-22, X, Y and M were extracted and sorted by coordinate. Overlaps between alignments and extracted gene features were identified using bedtools v2.31.1:

~~~
bedtools intersect -a {bam} -b {gtf} -wo -sorted -bed > {bed}
~~~

Each read was assigned to its longest-overlapping gene, recorded in the XT tag, with the XS tag set to “Assigned.” Reads which did not overlap a gene had the XS tag set to “Unassigned_NoFeatures” value given. The original demultiplexed barcode and UMI were retained in the UB and CB tag. UMI-tools v1.1.6 was run using Python v.3.9.21:

~~~
umi_tools dedup \
  --stdin={input_bam} --stdout={output_bam} \
  --temp-dir=“$TMPDIR” \
  --extract-umi-method=tag \
  --umi-tag=UB --cell-tag=CB \
  --per-cell --per-gene \
  --gene-tag=XT \
  --assigned-status-tag=XS \
  --no-sort-output
~~~

umi_tools group was also run with the same parameters to determine whether each deduplicated read belonged to a singleton or duplicate group. It was excluded from the runtime and memory benchmarks.

### Assessing consensus quality

In order to assess consensus quality, the duplicate group representative reads were aligned against the reference genome (GRCm38.p4 for mouse and GRCh38.p13 for human samples) using minimap2 v2.28-r1209 with parameters -ax splice -y --secondary=no -c --MD --eqx. The error rate was computed as (X + I + D) / (= + X + I), where X is the number of substitutions, I is the number of inserted bases, D is the number of deleted bases and = is the number of bases matching the reference genome based on the alignment CIGAR string.

Benchmarks were orchestrated using Snakemake v9.9.0. The pipeline, along with all preprocessing and figure generation code, is available at https://github.com/davidsongroup/nailpolish_benchmark.

### Assessing Nailpolish performance

Time and memory requirements were recorded on Linux high-performance nodes with Intel(R) Xeon(R) Gold 6342 CPUs. For Nailpolish, we report the combined runtime of indexation and consensus calling. We excluded any preprocessing required for UMI-tools and SiCeLoRe, including alignment, as Nailpolish does not require reads to be mapped beforehand.

### SNV calling

Reads from pre- and post-deduplication were first aligned with minimap2 v2.28 (−ax splice:hq -y --MD) using the reference from the LongBench GRCh38 assembly (GRCh38.primary_assembly.genome.add.spikein.fa), comprising the primary chromosomes, unplaced GL/KI scaffolds and synthetic spike-in contigs, then coordinate-sorted and indexed with samtools v1.23.1. Small variants were called with Clair3-RNA v0.2.2 using the ont_r10_dorado_cdna platform model, with --include_all_ctgs, --enable_phasing_model and otherwise default parameters; the phasing-model output was taken as the call set. Analyses were restricted to biallelic PASS records on chr1-22, X and Y. Recovery was scored against the Sanger whole-exome CaVEMan call set for SIDM00698 (Cell Model Passports; https://cellmodelpassports.sanger.ac.uk/passports/SIDM00698): 2,414 PASS single-nucleotide variants on GRCh38 with chr-prefixed contig names, normalised with bcftools v1.23 (norm -c w; zero reference-allele mismatches). Two subsets of this call set were scored separately: the 1,000 variants annotated as exonic against merged GENCODE v49 exons and the 175 variants sequenced to at least 10× in both pre- and post-deduplicated reads. Per-site depth at every truth position was measured per arm with samtools depth -a -b.

### Fusion detection

Fusions were called with JAFFAL v2.5 using its pre-built reference (GENCODE v49) with default parameters and identical settings in both pre- and post-deduplication. Recovery was scored against an independent RNA-level call set: JAFFAL calls from ONT dRNA of the same H146 cell line (https://www.biorxiv.org/content/biorxiv/early/2026/06/10/2025.09.11.675724/DC6/embed/media-6.xlsx?download=true; Supplementary Table 5 from You *et al*.), which is independent of the treatment evaluated here in that it is a different library preparation and no deduplication was applied to it. True fusion pairs were those carrying at least three spanning reads in the direct-RNA data (54 pairs); a call was scored as recovered when both partner genes matched a truth pair in either orientation.

## Abbreviations

UMI: Unique Molecular Identifier
ONT: Oxford Nanopore Technologies
FDD: False duplicate detection

## Declarations

### Availability of data and materials

Nailpolish is available from https://github.com/davidsongroup/nailpolish. Code to reproduce the figures in this paper is available at https://github.com/davidsongroup/nailpolish_benchmark. Raw datasets used for benchmarking are available from the Sequence Read Archive (SRA) under accessions PRJNA341465, PRJNA713904, PRJNA1171488, PRJNA644362, PRJNA1332620 and from the

European Nucleotide Archive (ENA) under accessions PRJEB54718 and PRJEB28878. The Sanger whole-exome CaVEMan call set for H146 is available from Cell Model Passports under accession ID SIDM00698.

### Authors’ contributions

NMD and OC developed the concept. OC implemented Nailpolish and performed validation and benchmarking. JWT performed variant and fusion data analysis. NMD supervised and funded the research. OC and NMD wrote the manuscript. All authors redrafted and approved the manuscript.

## Acknowledgements

This research was undertaken with the assistance of the Milton HPC, supported by WEHI Research Computing. We thank Noorul Amin, Feng Yan, Yupei You, Shian Su and Hannah Coughlan for providing data for testing and for giving valuable feedback on Nailpolish. Anthropic’s Claude models assisted with manuscript language editing and code generation. All LLM-generated code was reviewed, tested, and verified by the authors, who take full responsibility for its correctness. We acknowledge the Wurundjeri people of the Kulin nation as the traditional owners and guardians of the land on which the work was performed.

