## Supplementary Figures and Tables for "Nailpolish: Reference-free UMI deduplication and consensus read generation"

**Supplementary Material:**

Supplementary Tables 1-2

Supplementary Figures 1-4

**Supplementary Tables**

| Tool | Read type | Library preparation | Required input for deduplication | Reference-free | Consensus |
|---|---|---|---|---|---|
| UMIErrorCorrect | Short | UMI-tagged reads | .bam generated by upstream pipeline step | No | Yes (position-wise or MAFFT) |
| FGBio GroupReadsByUmi | Short | UMI-tagged reads | .bam with UMIs in tag | No | Yes (position-wise) |
| UMIc | Short | UMI-tagged reads | .fastq with UMIs encoded in read | Yes | Yes (position-wise) |
| PRESTo BuildConsensus | Short | UMI-tagged reads (designed for lymphocytic repertoire) | .fastq with custom header format | Yes | Yes (position-wise) |
| longread_umi | Long | Targeted amplicon (dual-end UMI flanked by primers) | Individual .fastq per UMI group | Optional | Yes (POA) |
| umi-pipeline-nf | Long | Targeted amplicon (dual-end UMI flanked by primers) | Individual .fastq per UMI group | No | Yes (POA) |
| SiCeLoRe | Long | UMI-tagged reads | .bam with UMIs in tag | No | Yes (POA) for groups of size ≥ 3 |
| scNanoGPS | Long | UMI-tagged reads | .fastq with custom header format | No | Yes (POA) |
| isoseq | HiFi long reads | UMI-tagged reads; requires Kinnex library prep and HiFi sequencing | FLTNC .bam with custom tags | Yes | Yes (POA) |
| UMITools | Short and long | UMI-tagged reads | Sorted .bam with UMIs encoded in read or in the tag | No | No |
| **nailpolish** | **Short and long** | **UMI-tagged reads** | **.fastq with UMIs in a user-defined header or in a separate table** | **Yes** | **Yes (POA)** |

**Supplementary Table 1.** Comparison of tools for UMI-based deduplication and consensus calling. Tools are grouped by their primary target read type (short- or long- read) and sequencing protocol. Standalone indicates whether a tool is bundled independently or as part of a broader pipeline. Reference-free indicates that the tool does not require alignment prior to deduplication. Tools with consensus calling merge multiple reads to create a representative sequence, as opposed to selecting one. Position-wise consensus assumes that reads are co-terminal and bases at each position are chosen without an alignment step first.

| Dataset ID | Protocol | Description | Accession | Reads | Unique UMIs |
|---|---|---|---|---|---|
| scmixology2 | ONT PromethION R9.4.1/SQK-LSK109; FLT-seq | Equal mixture of cells from 5 cell lines (H2228, H838, H1975, HCC827, A549) | PRJNA341465 | 21822958 | 18650938 |
| GridION | ONT GridION R9.4.1/SQK-LSK110; FLT-seq | ~1000 human induced pluripotent stem cells (hiPSC)-derived neural progenitors | PRJEB54718 | 6617007 | 6328318 |
| GridION Q20 | ONT GridION R10.4/SQK-LSK110 and Q20+ kit; FLT-seq | | | 3030565 | 2876521 |
| PromethION | ONT PromethION R9.4.1/SQK-LSK110; FLT-seq | | | 53157076 | 42331289 |
| LR-split-seq | PacBio Sequel II (CLR) | Split-seq combinatorial barcoding of the C2C12 myogenic system in Mus muscularis | PRJNA713904 | 2346583 | 2081377 |
| LongBench SC ONT | ONT PromethION R10.4.1/SQK-LSK114; 10x Chromium v3 | Pooled mixture of cells from 8 cell lines (lung adenocarcinoma; small cell lung cancer (SCLC) molecular subtypes A and P) | PRJNA1171488 | 21737936 | 19306372 |
| LongBench SC PB | PacBio Revio & Kinnex; 10x Chromium v3 | | | 38591908 | 32784026 |
| LongBench bulk H146 | ONT PromethION bulk PCR-cDNA; SQK-PCB114 | H146 cell line (SCLC-A) | | 27842725 | 17548310 |
| Visium | ONT PromethION R9.4.1/SQK-LSK110; 10x Visium | Mouse brain olfactory bulb | PRJNA644362 | 26168349 | 12475461 |
| DNM1L | ONT PromethION R10.4/SQK-NBD114 | Targeted long-read sequencing of DNM1L | PRJNA1332620 | 2356402 | 507926 |
| RAGE-seq | ONT GridION R9.4.1/SQK-LSK108; 10x Chromium v2 | Targeted T- and B-cell long-read single cell sequencing of breast cancer primary tumour | PRJEB28878 (sample ERR3273784) | 6262648 | 3305522 |

**Supplementary Table 2.** Details of the RNA-seq datasets used.

#### Supplementary Figures

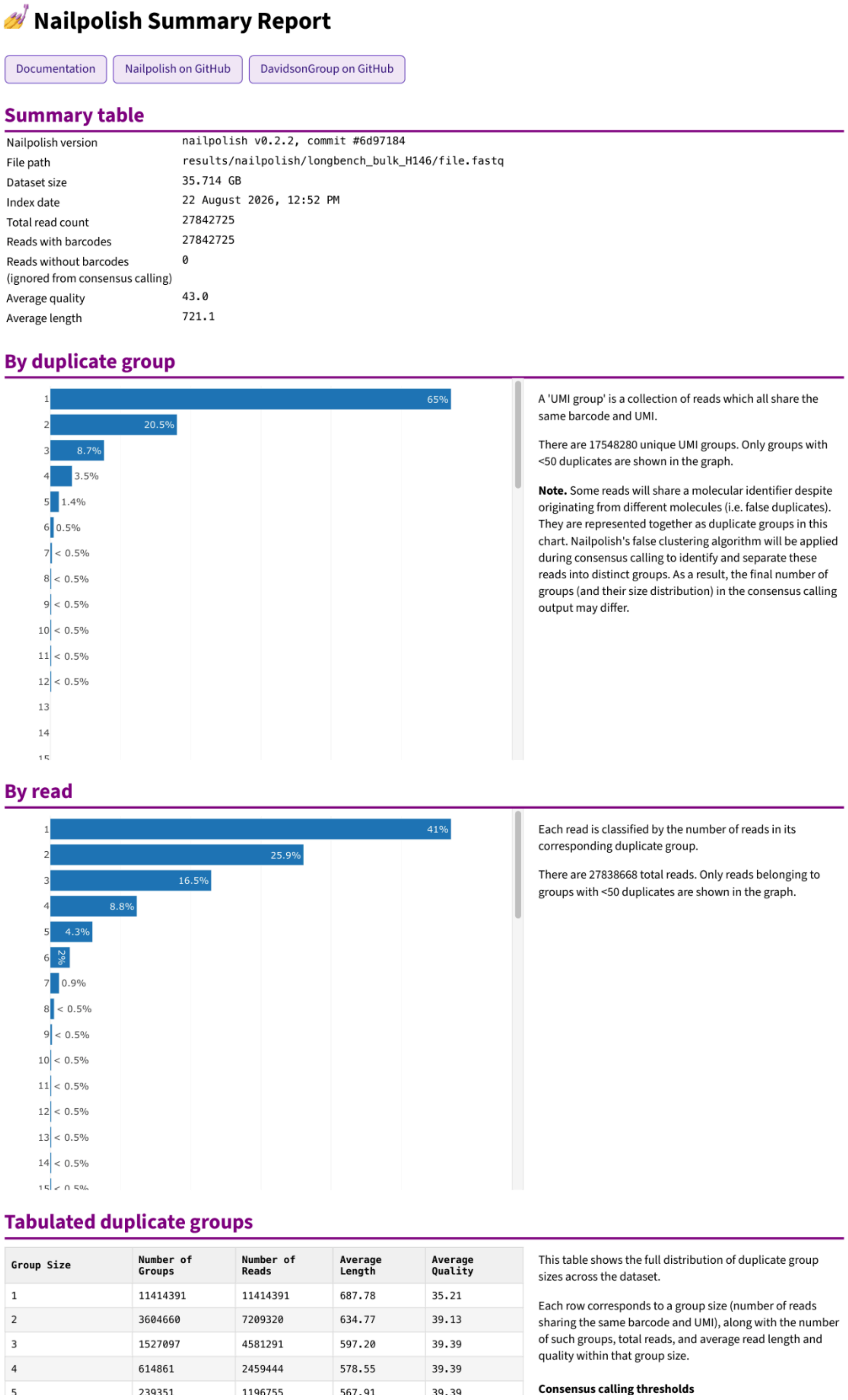

### Nailpolish Summary Report

Documentation | Nailpolish on GitHub | DavidsonGroup on GitHub

#### Summary table

| | |
|---|---|
| Nailpolish version | nailpolish v0.2.2, commit #6d97184 |
| File path | results/nailpolish/longbench_bulk_H146/file.fastq |
| Dataset size | 35.714 GB |
| Index date | 22 August 2026, 12:52 PM |
| Total read count | 27842725 |
| Reads with barcodes | 27842725 |
| Reads without barcodes (ignored from consensus calling) | 0 |
| Average quality | 43.0 |
| Average length | 721.1 |

#### By duplicate group



A 'UMI group' is a collection of reads which all share the same barcode and UMI.

There are 17548280 unique UMI groups. Only groups with <50 duplicates are shown in the graph.

**Note.** Some reads will share a molecular identifier despite originating from different molecules (i.e. false duplicates). They are represented together as duplicate groups in this chart. Nailpolish's false clustering algorithm will be applied during consensus calling to identify and separate these reads into distinct groups. As a result, the final number of groups (and their size distribution) in the consensus calling output may differ.

#### By read



Each read is classified by the number of reads in its corresponding duplicate group.

There are 27838668 total reads. Only reads belonging to groups with <50 duplicates are shown in the graph.

#### Tabulated duplicate groups

| Group Size | Number of Groups | Number of Reads | Average Length | Average Quality |
|---|---|---|---|---|
| 1 | 11414391 | 11414391 | 687.78 | 35.21 |
| 2 | 3604660 | 7209320 | 634.77 | 39.13 |
| 3 | 1527097 | 4581291 | 597.20 | 39.39 |
| 4 | 614861 | 2459444 | 578.55 | 39.39 |
| 5 | 239351 | 1196755 | 567.91 | 39.39 |

This table shows the full distribution of duplicate group sizes across the dataset.

Each row corresponds to a group size (number of reads sharing the same barcode and UMI), along with the number of such groups, total reads, and average read length and quality within that group size.

**Consensus calling thresholds**

**Supplementary Figure 1:** Example HTML summary report for the LB Bulk H146 dataset, generated by nailpolish summary following indexation. The report includes a summary table, including the read count, quality, and length. The distribution of duplicate group sizes are shown in graph form, both as a proportion of groups ("By duplicate group") and as a proportion of reads ("By read"), and as a tabular breakdown by group size. The page has been truncated for display.

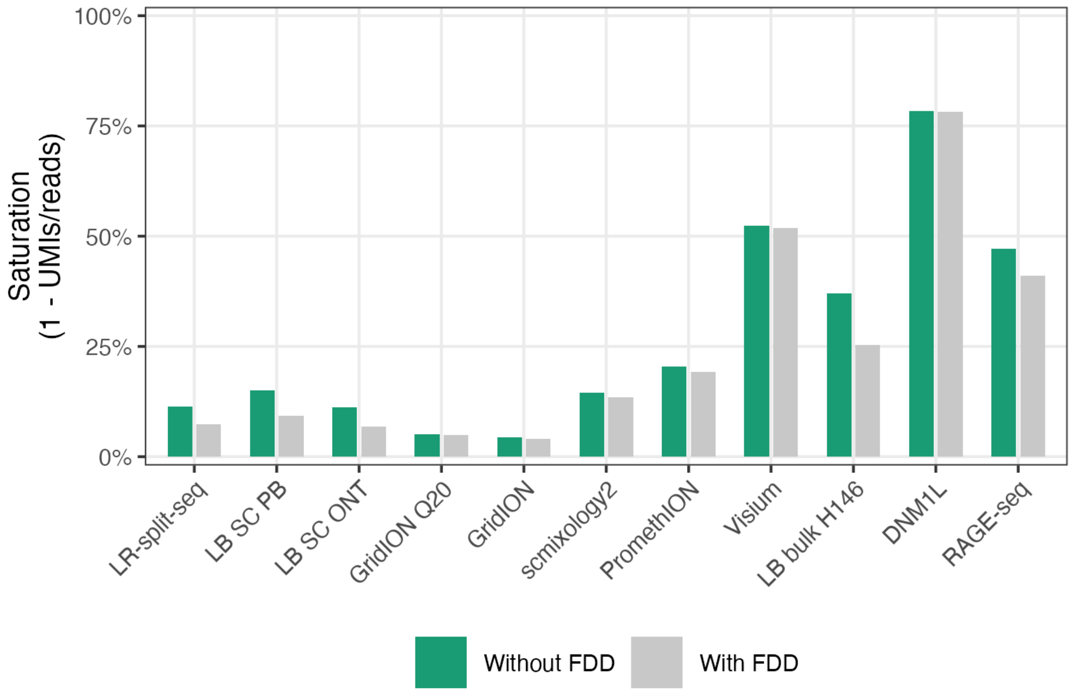


**Supplementary Figure 2:** Estimated sequencing saturation with and without false duplicate detection, where saturation was computed as 1 - ( number of unique duplicate groups / total reads ).

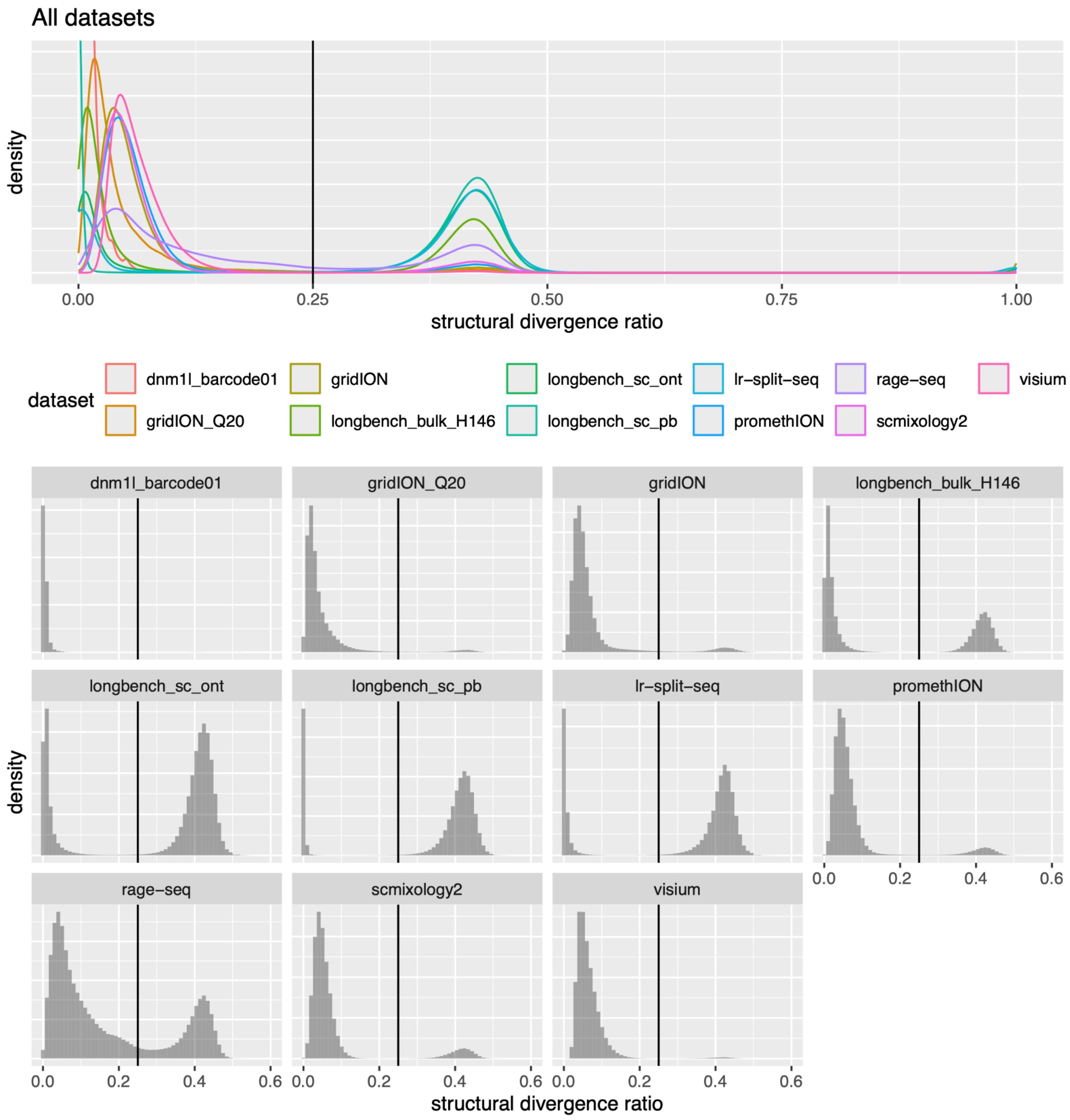


**Supplementary Figure 3:** Distribution of the structural divergence ratio across datasets. Top: the density of all 11 datasets overlaid. Bottom: histograms per dataset, with y-axes scaled independently. In all figures, the vertical line marks the default threshold of 0.25, above which candidate reads are classified as false duplicates.

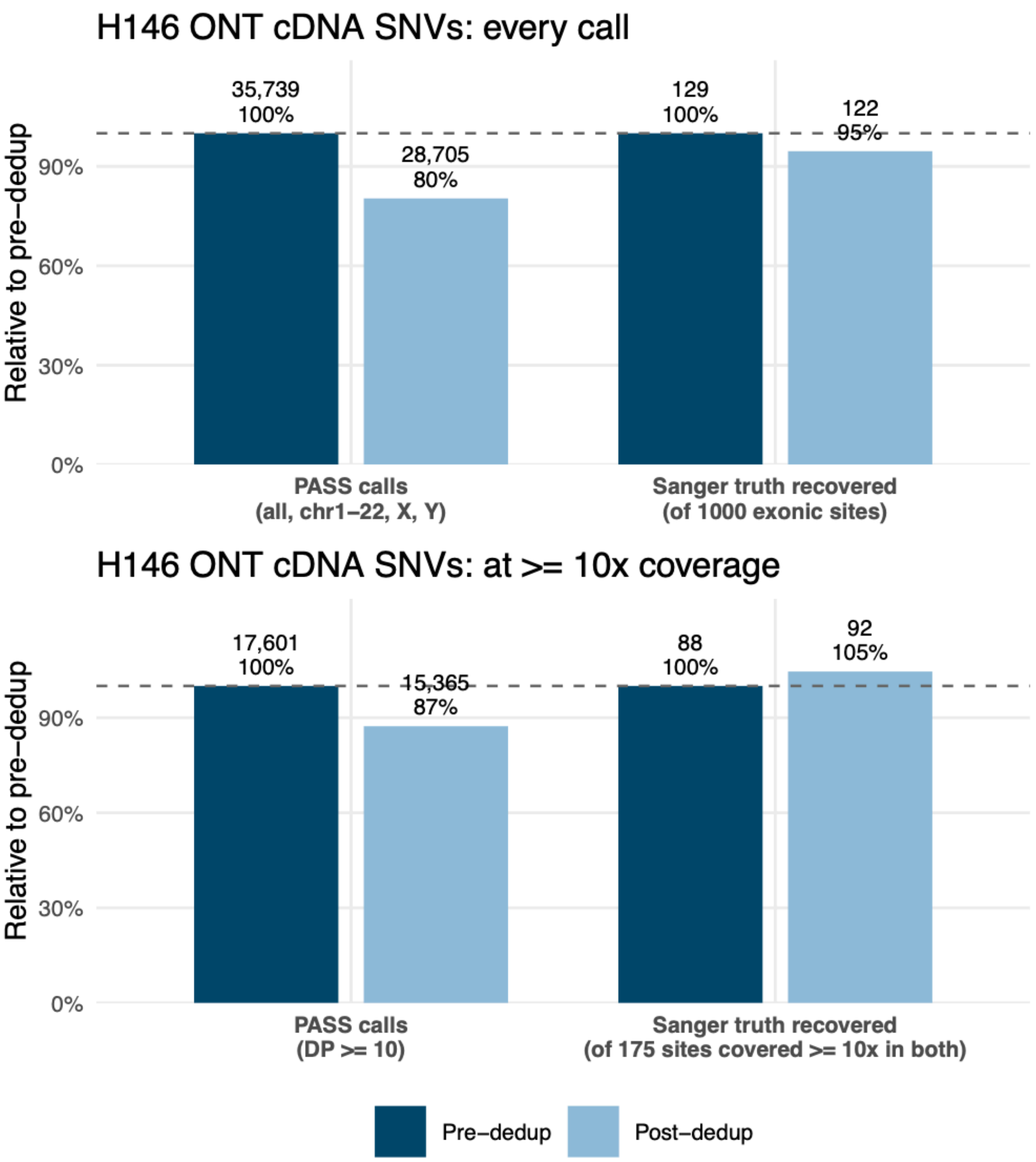


**Supplementary Figure 4:** Clair3-RNA calls from the H146 bulk ONT cDNA sample before and after Nailpolish deduplication: (Top) All biallelic PASS calls on chr1-22, X and Y (left), against recovery of the 1,000 exonic Sanger truth variants (right). (Bottom) Calls at read depth ≥10 (left), against recovery of the 175 truth variants sequenced to ≥10× in both arms (right).
